# Human Neural Synergy when combining Stevia with a Flavor Modifer and the Neural effects of Sucrose vs Stevia

**DOI:** 10.1101/2025.06.17.657228

**Authors:** Hee-kyoung Ko, Jingang Shi, Thomas Eidenberger, Weiyao Shi, Ciara McCabe

## Abstract

There is a drive to improve the acceptability of sweeteners like stevia by reducing their off-tastes. The main aim was to examine the synergistic neural effects of combining stevia with a flavor modifier and secondly to examine stevia vs. sucrose due to limited human neuroimaging data and concerns that sweeteners may be more addictive than sugar.

In a within-subjects fMRI study, 34 healthy adults (Mean age = 25) tasted four conditions: stevia, stevia plus a flavor modifier, the modifier alone, and sucrose. We analyzed whole-brain responses and focused on regions of interest (ROIs) including the insula, postcentral gyrus, and hypothalamus (identified via meta-analysis of sweet taste processing), as well as the nucleus accumbens (NAcc) and amygdala due to their roles in reward and aversion.

Stevia combined with the modifier evoked super-additive responses in the postcentral gyrus, parietal cortex, and occipital gyrus (p < 0.05, FWE-corrected). Compared to stevia alone, the stevia-modifier combination elicited reduced hypothalamic activity (p = 0.008) and the hypothalamus tracked pleasantness and mouth fullness only in this condition. The NAcc tracked mouth fullness more for the modified stevia than for stevia alone, and the amygdala tracked bitterness only in the plain stevia condition. Sucrose elicited higher postcentral gyrus activation than stevia (p = 0.01).

We provide first evidence that combining stevia with a flavour modifier reveals synergistic neural activity associated with taste sensation, intensity and multisensory integration. Adding a modifier to stevia could increase unconscious desirability for stevia by masking its bitterness and increasing its mouth fullness.

## Introduction

Excessive intake of free sugars threatens the nutrient quality of food and leads to unhealthy weight gain and an increased risk of being overweight or obese, according to the World Health Organisation report (WHO, 2023).

Using high intensity non-nutrient sweeteners or non-caloric/low_-_calorie sweeteners — such as saccharin, sucralose, aspartame, cyclamate, and acesulfame_-_K in food and beverages, can help reduce sugar (Shankar et al., 2013) and lead to weight reduction (Rogers & Appleton, 2021) particularly in those overweight/obese under an unrestricted diet (Laviada_-_Molina et al., 2020). Studies find that sugar substitutes such as stevia a natural zero-calorie sweetener, has advantageous effects on appetite and energy intake (Stamataki, Crooks, et al., 2020; Stamataki, Scott, et al., 2020). While individuals are encouraged to make healthier choices to reduce sugar intake, governments also recognize the vital responsibility of the food industry in promoting healthier diets, particularly through efforts to lower sugar levels in processed foods (Miller et al., 2021).

To do this food companies can replace sugar with non-nutrient sweeteners such as stevia. Sugar-like alternatives should be developed that replicate both the pleasantness and mouthfeel of sugar, without relying on high concentrations of sweeteners. One approach involves using flavorings with modifying properties (FMPs), which can enhance or mask a food’s sensory profile without introducing new flavor characteristics (EU, 2014). This could be particularly useful for natural sweeteners like stevia that lack mouth fullness and have prolonged sweet lingering, herbal notes and bitterness at higher concentrations (Ashwell, 2015; Muenprasitivej et al., 2022; Peteliuk et al., 2021).

Our previous study (Ko et al., 2025), the first of its kind, found that by adding a FMP to the sweetener sucralose resulted in super-additive neural effects in regions of the brain related to taste sensation and multisensory integration. This is consistent with prior neuroimaging studies showing super-additive neural effects of combining sucrose and sweet odors (De Araujo et al., 2003; Small et al., 2004). Specifically, we found synergistic activity in the mid/inferior temporal gyri, pre and postcentral gyri and parietal areas, which could underpin potential mechanisms by which non-nutrient sweeteners could be made more acceptable in diets, thus aiding sugar replacement (Ko et al., 2025).

Therefore, the main aim of this study was to examine for the first time the neural synergistic effects of adding a FMP to the natural sweetener stevia. To do this we used a FMP Proust 221 (EPC Natural Products Co.) evaluated for safety and compliance with the guidelines established by the Flavor and Extract Manufacturers Association of the United States (Harman & Hallagan, 2013). As Proust 221 is designed to enhance sweetness this also allows us to examine neural effects outside of any perceptual differences in sweetness between when modifier is and isn’t added to the stevia. We hypothesized that by adding the FMP Proust 221 to stevia we could augment its sensory profile, enhancing its neural similarity to sucrose.

How brain activity to the taste of stevia might differ from sucrose is unclear, we know of no studies directly comparing these tastes using fMRI. Connolly et al. (2013) reported that a network of brain regions—including the anterior insula, right anterior cingulate cortex, bilateral amygdala, left hippocampus, and visual cortex—remained similarly active up to 30 minutes after consumption of sucrose- and stevia-flavored drinks. In a subset of obese participants (N = 10), there was some indication of greater activation in response to sucrose compared to stevia, although this was not based on a direct head-to-head comparison (Connolly et al., 2013). Another study reported greater fluctuations in resting-state neural activity 10 minutes after sucrose ingestion compared to stevia, specifically in the nucleus tractus solitarius—a region involved in sensory integration—in both obese (N = 11) and lean (N = 11) women (Kilpatrick et al., 2014). The authors proposed that these neural differences might reflect distinct nutrient sensing mechanisms. However, a more recent meta-analysis examining the effects of sweet tastes on the human brain found no consistent differences in neural activity between sucrose and non-nutritive sweeteners, including stevia, even within reward-related regions. The meta-analysis also noted that the limited number of studies in this area often suffer from small sample sizes and overly liberal statistical thresholds (Yeung & Wong, 2020). This is somewhat consistent with our recent finding of no significant differences between the sucrose and sucralose at the whole brain or in reward related regions but greater somatosensory activity to sucrose vs sucralose in the postcentral gyrus ROI (Ko et al., 2025).

Therefore, our second study aim was to directly examine the neural response to sucrose vs stevia by delivering the tastes to the mouths of participants whilst they were in the MRI machine. We aimed to recruit a robust sample size for fMRI (N=34) and employ a standardised fasting and feeding protocol. We hypothesised increased activity to sucrose vs stevia given our recent findings with sucralose. We also aimed to examine the relationships between neural activity and the subjective experiences of the tastes, pleasantness, bitterness, and mouth fullness across conditions. By clarifying the neural pathways responsible for these sensory experiences, this research could provide valuable insights for improving the sensory qualities of stevia-based products.

## Materials and Methods

### Participants

Thirty-four healthy and right-handed adults (10 male and 24 female) were recruited for the fMRI study. All participants were between 18 and 45 years old (mean 25 yrs) and had a current body mass index (BMI, weight in kg/height in m2) or waist-to-height ratio (WTH) in the healthy range. Participants were excluded if they had any current/previous psychiatric history using the Structured Clinical Interview for DSM-IV Axis I Disorder Schedule (SCID), or if they took psychoactive medication or an eating disorder (measured with Eating Attitude Test (EAT) > 20), food allergies, diabetes, smoking, or any contraindications to fMRI scanning. We also recorded the frequency, liking and craving for sugary and sweetened foods (Rolls & McCabe, 2007). The questions in this scale consisted of “How frequently do you eat sugary foods?” with answers of either; a few times per month; 1-2 times per week; 3-4 times per week; or more than 5 times per week and “How frequently do you eat/drink foods with sweeteners?”, with answers of either; Never; Rarely; Sometimes; Often; Usually or Always. The Craving and Liking for sugary foods were scored as 1 for low craving and 10 for high craving on a Likert scale. All procedures contributing to this work comply with the ethical standards of the Helsinki Declaration of 1975, as revised in 2013 and ethical approval was obtained from the University of Reading Ethics committee, ethics ref: 2023-130-CM all participants provided written informed consent.

### Pre-test 1 (Triangle test or Taste perception test)

The 34 participants were entered into the study as long as they could distinguish 2% sucrose from a control. This standard taste perception test was as follows: The participants were randomly allocated to the following sequences of two samples A (distilled water) and B (20 g sucrose/litre [2 % Sucrose]): ABB, AAB, ABA, BBA, BAA and BAB. For the individual performance, each participant received all six sequences in random order. In a sequence, the participants took the whole 10 mL of each sample into their mouth, swirled and coat the solution around their mouth for 3 seconds and then spit it into a spittoon. On each trial after tasting all three, they indicated which was different from the other two. Participants who yielded correct identification of at least 5 out of the 6 trials on a second attempt, were recruited to the study.

### Stimuli for the scan

Stevia (ST), the flavour modifier Proust 221 (M) the combination of stevia with flavour modifier Proust (STM) and Sucrose (S) provided the basic stimuli set for the fMRI study. The sucrose used was >99.7% pure with less than 0.04 % inverted sugar (i.e. fructose and glucose) and less than 0.06 % loss during drying, and sourced from Wiener Zucker, Feinkristallzucker, Austria and the sweet concentration of sucrose was 6% (Wee et al., 2018). The stevia sample consisted of ≥ 95 % steviol glycosides, thereof ≥ 97 % Rebaudioside A. The flavour modifier (Proust 221, 0.036%) was provided by EPC Natural Products Co., Ltd. As we wanted to keep the stimuli perceptually similar we matched the stevia concentration (0.036%) to an equally sweet concentration of sucrose 6% (Wee et al., 2018) and reduced the stevia (0.032%) when combining them with the modifier, as the modifier can increase sweetness (Wee et al., 2018). A tasteless solution (containing the main ionic components of saliva, 25 mM KCl + 2.5 mM NaHCO3) was used as a control. The sucrose and stevia were diluted and delivered in 100ml distilled water, as we wanted to keep the modifier tasteless it was diluted and delivered in 100ml control tasteless solution (containing the main ionic components of saliva, 25 mM KCl + 2.5 mM NaHCO3). The control tasteless solution was also used as a control rinse between trials (Table 1).

**Table 1.**

| **Table 1** |  | Solution |
| --- | --- | --- |
| Stevia + Modifier | STM | 0.032 g stevia (Reb-A, Steviolglycoside) and 0.004 mg Proust 221 |
| Stevia | ST | 0.036 g stevia (Reb-A, Steviolglycoside) |
| Modifier | M | 0.04 g Proust 221 |
| Sucrose | S | 6 g sucrose |
| Control | C | 25mM KCl + 2.5 mM NaHCO<sub>3</sub> |

The two-alternative forced choice (2AFC) sensory tests were done to verify that the sweetness perception of the low stevia concentration plus modifier was equal to the higher concentration of stevia alone. To confirm that the modifier does not have a sweet taste of its own we also compared it to 1.5% sucrose, the lowest known concentration of a sweet taste perceivable by humans (Petty et al., 2020). Twelve expert sensory panelists recruited from the University of Applied Sciences Upper Austria, were given three times in a random blinded order a pair of samples and asked to decide which of the samples tasted more/less sweet. In total 12 x 3 = 36 ratings were obtained for each analysis and were statistically evaluated using the beta-binomial evaluation.

### Study design

The fMRI scans took place at the Centre of Integrative Neuroscience and Neurodynamics (CINN) at the University of Reading. If scheduled for a morning scan participants fasted overnight, if having an afternoon scan participants fasted for 3 h (no food, only water) before the scan. 10 participants had a morning scan, and 24 participants had an afternoon scan. 60-90 minutes before scanning all the participants were given a standardized meal similar to previous studies (a banana, a cup of orange juice, 2 crackers, ∼261 total calories) with the instruction to “eat until feeling comfortably full, without overeating” similar to our previous study (Thomas et al., 2015). We asked participants to rate their hunger and mood, before the scan, on a visual analogue scale from 0 being not at all to 10 indicating the most ever felt. Subjects were screened for potential pregnancy and metal in their body before being placed in the fMRI scanner.

The task consisted of a block design where we had 7 trials each of stevia (ST), stevia & modifier (STM), sucrose (S), and modifier (M) in a pseudo randomised order. There was also a break (∼7 min) after any blocks that included ST to allow for any lingering effects of ST to dissipate. Tastes were delivered to the subject via separate long (∼3 m) thin Teflon tubes with a mouthpiece (∼1cm in diameter) at one end, which was held by the subject comfortably between the centre of the lips. At the other end of the tubes were connected to separate reservoirs via syringes and one-way Syringe Activated Dual Check Valves (Model 14044– 5, World Precision Instruments, Inc) which allowed any stimulus to be delivered manually by the researcher at exactly the right time indicated by the programme (Murray et al., 2014) thus avoiding the delays and technical issues experienced when using computerised syringe drivers.

### fMRI Task

At the beginning of a trial, a white cross at the centre of the screen appeared for 2 s indicating the start. Then, a stimulus was delivered in a 0.5 mL aliquot to the subject’s mouth, the green cross was presented at the same time on the visual display for 5 s. The instruction given to the subject was to move the tongue once as soon as the stimulus was delivered in order to distribute the solution round the mouth to activate receptors, and then to keep still until a red cross was shown, when the subject could swallow. Swallowing was 2 s, then the subject was asked to rate the ‘pleasantness’ (+2 to –2), the ‘bitterness’ (0 to +4), and the ‘mouth fullness’ (richness) (0 to +4) of the taste in their mouth, measuring hedonic and sensory aspects of the stimulus, on a visual analogue scale by moving a bar to the appropriate point on the scale using a button box, ratings similar to those used in previous taste/fmri studies (Rolls, 2019). Each rating period was 5 s long. After the last rating on each trial, 0.5mL of the control (tasteless) solution was administered in the same way as the experimental stimulus and a green cross was again presented at the same time on the visual display for 5 s. The control was used as the comparison condition to allow somatosensory effects produced by liquid in the mouth, and the single tongue movement made to distribute the liquid throughout the mouth, to be subtracted in analysis (De Araujo et al., 2003; O’Doherty et al., 2001). The tasteless control condition was not subjectively rated. After the control rinse stimulus a grey cross was presented for a duration between 0.8 s and 2 s (jittered) to indicate the end of the trial. Then the screen was black for 2 s before a new trial started. Each trial lasted ∼30 sec.

### fMRI data acquisition

Blood oxygenation level dependent (BOLD) functional MRI images were acquired using a three-Tesla Siemens scanner (Siemens AG, Erlangen, Germany) with a 32-channel head coil. During the task, 730 volumes were obtained for each participant, 54 slices, using a multiband sequence with GRAPPA and an acceleration factor of 6. Other sequence parameters included a repetition time (TR) of 700ms, an echo time (TE) of 30ms, and a flip angle (FA) of 90°. The field of view (FOV) covered the whole brain with a voxel resolution of 2.4 x 2.4 x 2.4 mm^3^. Structural T1-weighted images were acquired utilizing a magnetization prepared rapid acquisition gradient echo sequence (TR = 2300ms, TE = 2.29ms, FA = 8°) with a FOV covering the whole brain and a voxel resolution of 1 x 1 x 1mm^3^.

### fMRI data analysis

The imaging data were analysed using SPM12 (Wellcome Centre for Human Neuroimaging, University College London). Pre-processing of the data used SPM12 realignment, coregister, segment, normalization to the MNI coordinate system (Montreal Neurological Institute; Collins et al., 1994), and spatial smoothing with a 6 mm full width at half maximum isotropic Gaussian kernel. The time series at each voxel was low pass filtered with a haemodynamic response kernel. Time series non-sphericity at each voxel was estimated and corrected for, with a high-pass filter with cut-off period of 128 s.

In the single-event design, a general linear model was then applied to the time course of activation in which stimulus onsets were modelled as single impulse response functions and then convolved with the canonical hemodynamic response function. Linear contrasts were defined to test specific effects. Following smoothness estimation, linear contrasts of parameter estimates were defined to test the specific effects of each condition with each individual data-set. Voxel values for each contrast resulted in a statistical parametric map of the corresponding t statistic (transformed into the unit normal distribution (SPM z)). Movement parameters and were added as additional regressors.

We then examined if areas were more strongly activated by the combination stevia plus modifier than by the sum of any activations produced by the two stimuli presented separately e.g. STM> Sum (ST, M) in the whole brain, similar to our previous studies (Ko et al., 2025; McCabe & Rolls, 2007). We also examined the direct comparison of stevia plus modifier vs stevia alone and sucrose vs stevia in the whole brain. We thresholded at p<0.05 corrected (familywise-error (FWE) and p values cluster corrected at both p<0.05 False Discovery Rate (FDR) and p<0.05 FWE. We also checked if the results were affected by gender, hunger level and scan time (morning or afternoon) added as covariates of no interest.

We then examined five regions of interest (ROI), the postcentral gyrus, insula, hypothalamus, nucleus accumbens and amygdala. Spheres (10mm) were created for the anterior insula (primary taste cortex, [-32, 16, 2]), posterior insula [-38,-2,-12], and postcentral gyrus [60 −16 24] using WFU pickatlas, and the hypothalamus using aal atlas, identified from meta-analyses on sweet tastes in humans (Roberts et al., 2020; Yeung & Wong, 2020). Given we are interested in the possible rewarding and aversive effects of stevia we also examined the nucleus accumbens (Berridge, 2009) and amygdala (Gottfried et al., 2003; Izadi & Radahmadi, 2022) using (IBASPM71 atlas) and aal atlas anatomical masks, respectively, in WFU pickatlas. For the ROI analyses, data were extracted using the SPM ROI analysis Matlab code and SPM’s spm_get_data command and analysed with paired-sample t tests, in excel and SPSS, and then corrected for multiple comparisons across the 5 ROIs, i.e. p=0.05/5=0.01.

We also did exploratory analysis examining the relationship between whole brain neural activity and the subjective ratings (pleasantness, bitterness and mouth fullness) across all participants using a multiple regression analysis in SPM12. As there was very little variability in the subjective ratings, (as we designed the study to make it difficult for participants to distinguish between stimuli) we used a lenient threshold of p=0.05 uncorrected. For example, all participants’ scans for the condition stevia were entered into a model as a regressor with the corresponding participant’s subjective pleasantness ratings added as an additional regressor. This allowed us to run correlations between neural activity and ratings.

## Results

### Sensory Results

Examining data from twelve sensory panelists, (mean 23 yrs, SD, 3.78) 7 female and 5 male with healthy weight (BMI mean 22.1, SD, 4.66) we found no statistical difference in sweetness equivalence of the conditions Stevia (Reb-A, Steviolglycoside 0.036 %) and sucrose (6 %) (Table 2).

**Table 2.** Sensory Data: Sweetness Equivalence (more/less sweet)

| Replicate 1 | Replicate 2 | Replicate 3 | Sum | p-value |
| --- | --- | --- | --- | --- |
| Reb-A (ST) 0.036 % vs Sucrose (S) 6% |  |  |  |  |
| 4/8 | 6/6 | 7/5 | 17/19 | 0.384 |
| Reb-A (ST) 0.032 % plus Proust 221 (M) vs. Reb-A (ST) 0.036 % |  |  |  |  |
| 7/5 | 5/7 | 8/4 | 20/16 | 0.351 |
| Proust 221 (M) vs. Sucrose (S) 1.5%, |  |  |  |  |
| 4/8 | 3/9 | 2/10 | 9/27 | 0.001 |
| In a random blinded order participants rated a pair of samples and were asked to decide which of the samples tasted more/less sweet. |  |  |  |  |

**Table 3:** Demographics.

| **Table 3: Demographics** | **N-34**<br>**Mean (SD)** |
| --- | --- |
| Age, years | 25.71 (8.25) |
| Gender, female/male: *n* | 24/10 |
| Body mass index | 22.00 (2.68) |
| EAT | 3.09 (3.20) |
| Craving for sugary foods | 5.11 (1.99) |
| Liking for sugary foods | 5.85 (1.98) |
| Freq eating sugary foods | 3.44 (2.09) |
| Freq eating/drinking foods with sweeteners | 3.97 (2.11) |

When examining the sweetness equivalence of the mixture of stevia (Reb-A, Steviolglycoside 0.032 %) plus the modifier Proust 221 vs. stevia alone (Reb-A, Steviolglycoside 0.036 %) the results show no statistical difference between the two samples for sweetness (Table 2).

When examining the sweetness equivalence of Proust 221 vs. sucrose 1.5%, Proust 221 was shown to be significantly less sweet than a 1.5 % sucrose solution which is considered the lowest perceivable concentration of a sweet taste for humans (Petty et al., 2020). Indicating that Proust on its own has no perceptible sweetness, as expected (Table 2).

### Pre-test results of sensitivity to 2% sucrose

Twenty-one participants passed the pre-test with 6 out of 6 trials correct the first time. Ten participants passed the pre-test with 5 out of 6 trials correct the first time and three participants got 6 of the 6 trials correct on their second attempt, so were also included in the study.

### fMRI scan day

#### Subjective hunger and Mood

Participants had relatively high mood and low hunger levels before the scan (Table 4).

**Table 4:** Visual Analogue Scale.

| **Table 4: Visual Analogue Scale** | **Mean score (± SD)** |
| --- | --- |
| **Appetite** |  |
| How hungry do you feel right now? | 4.35 ± 2.30 |
| How full do you feel right now? | 4.05 ± 2.11 |
| **Mood** |  |
| Alertness | 6.08 ± 2.40 |
| Disgust | 0.91 ± 1.23 |
| Drowsiness | 3.05 ± 2.66 |
| Anxiety | 1.79 ± 1.55 |
| Happiness | 6.11 ± 1.93 |
| Nausea | 0.70 ± 0.97 |
| Sadness | 0.55 ± 1.05 |
| Withdrawn | 1.08 ± 1.76 |
| Faint | 1.08 ± 1.84 |
| *Rate between 1 and 10, where 1 = Not at all, 10 = Most ever felt* |  |

#### Subjective ratings of stimuli

With a repeated measures ANOVA, with the subjective ratings as one factor with 3 levels (pleasantess, bitterness and mouth fullness) and condtion as the second factor (S, ST, STM). We found a main effect of ratings (F=19.7 (1.49,49) p<0.001), a main effect of condition (F=10 (2,66) p<0.001), and ratings * condition interaction (F=9 (4,132) p<0.001). Follow up paired sample t-tests found that there was a significantly increased rating of pleasantness for S vs STM (t=3.19 p=0.003) and S vs ST (t=6.53, p<0.001) and increased bitterness for ST vs S (t=2.69 p=0.01) and for STM vs S (t=2.62 p=0.01) and increased mouth fullness for S vs ST (t=3.41, p=0.002) see Figure 1.

**Figure 1.**
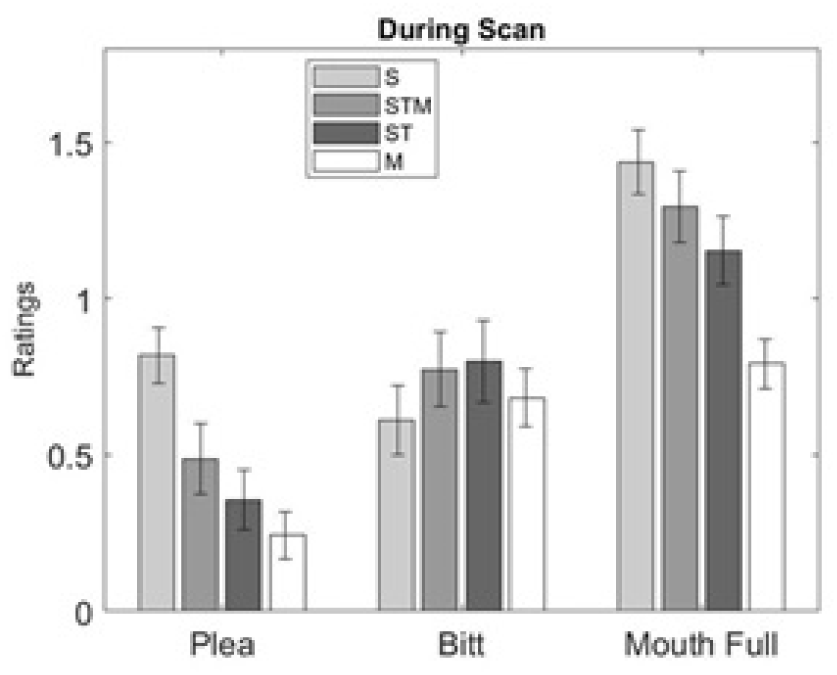
Subjective ratings during the scan.

### Whole brain analyses

#### Main effects of taste stimuli

As expected, similar to our previous studies the tastes in the mouth activated regions such as the primary taste cortex (insula), pre and postcentral gyrus, and the striatum (Table S1-S3). The sucrose-control condition also activated the mid cingulate and the hippocampus.

#### Super-additivity: STM > Sum (ST, M)

To examine any super-additive effects of combining the modifier with the stevia (STM) i.e., to examine if there was more activity to the combination than to the sum of the parts, even when sweetness was controlled, we used the contrast STM> Sum (ST, M). We found activity in regions similar to that of the individual tastes alone such as the pre and post central gyri (Table S1-S3). However, the parietal cortex and the occipital gyrus were also activated (Table 5).

**Table 5.**
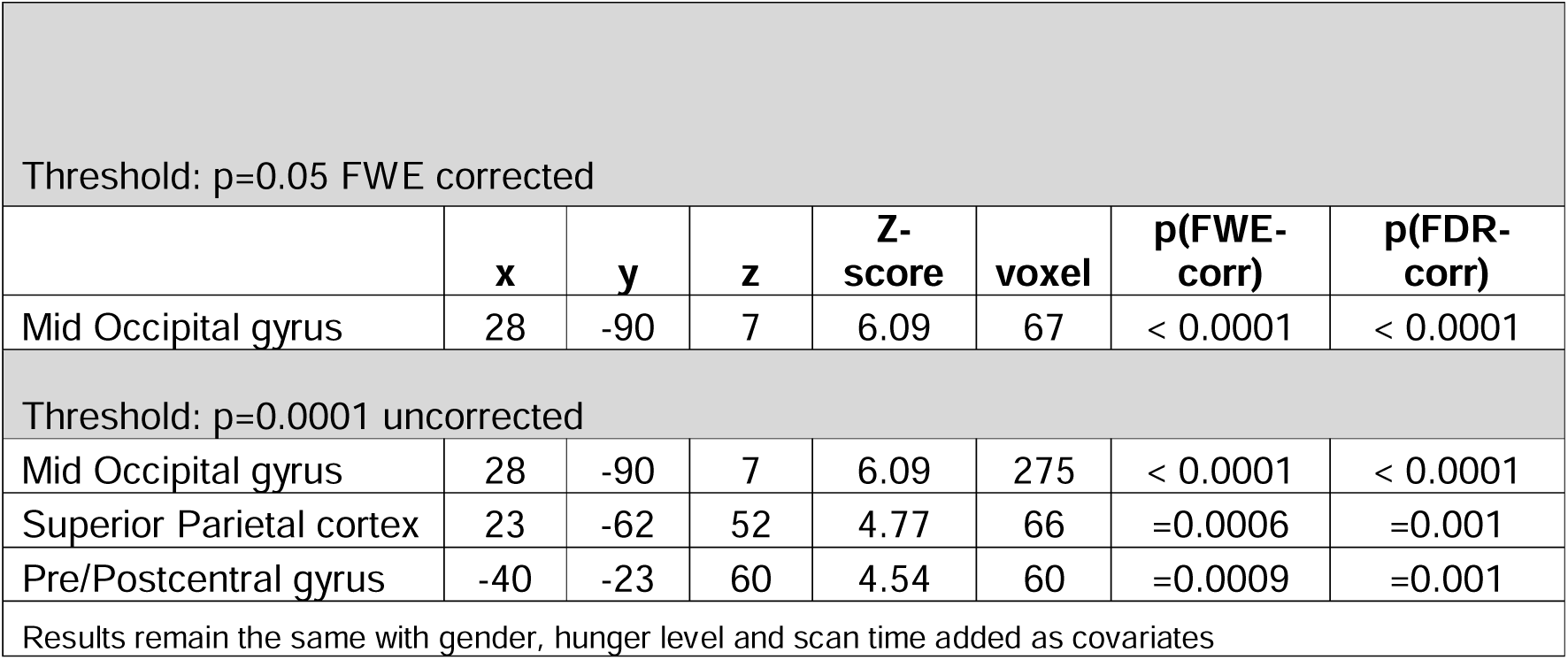
STM > Sum (ST, M)

#### STM vs ST

There were no whole brain differences between stevia plus modifier vs stevia conditions.

#### STM vs S

There were no whole brain differences between stevia plus modifier vs sucrose conditions.

#### S vs ST

When examining the whole brain results (Table 6) we found increased activity for the contrast S vs ST in the insula (figure 2) and rolandic operculum, these results were only apparent when using a p=0.001 uncorrected threshold. There were no regions that were increased under the contrast ST vs S, at any threshold.

**Figure 2.**
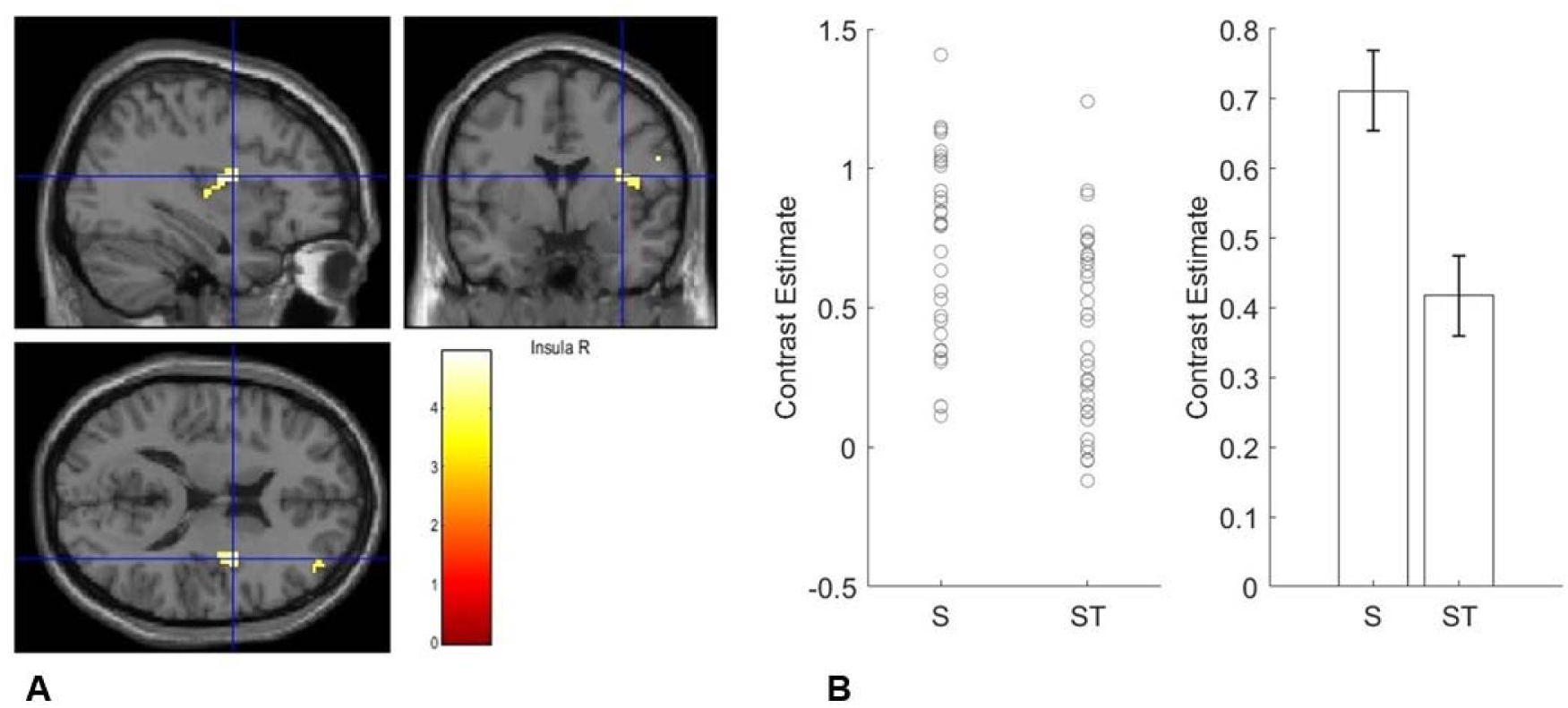
**A**. Sucrose (S) vs Stevia (ST) contrast **B**. Contrast estimates extracted from the insula for the S and ST conditions separately.

**Table 6.** Sucrose (S) – Stevia (ST)

Threshold: $p=0.001$ uncorrected
| Region | x(mm) | y(mm) | z(mm) | Z-score | Num of voxel | p(FWE-corr) | p(FDR-corr) |
| --- | --- | --- | --- | --- | --- | --- | --- |
| Insula | 35 | -2 | 16 | 4.25 | 61 | =0.038 | =0.052 |
| Rolandic operculum | 44 | 3 | 12 | 3.76 |  |  |  |

### ROI analyses

#### STM vs ST

We found decreased neural activity in the STM condition vs the ST condition in the hypothalamus ROI (t=2.52, p= 0.008, Cohens d =0.43) (Figure 3) and left anterior insula ROI (t=2.11, p= 0.02, Cohens d =0.36) (Table S4) (Figure 4). However, only the hypothalamus survived corrections for multiple comparisons.

**Figure 3.**
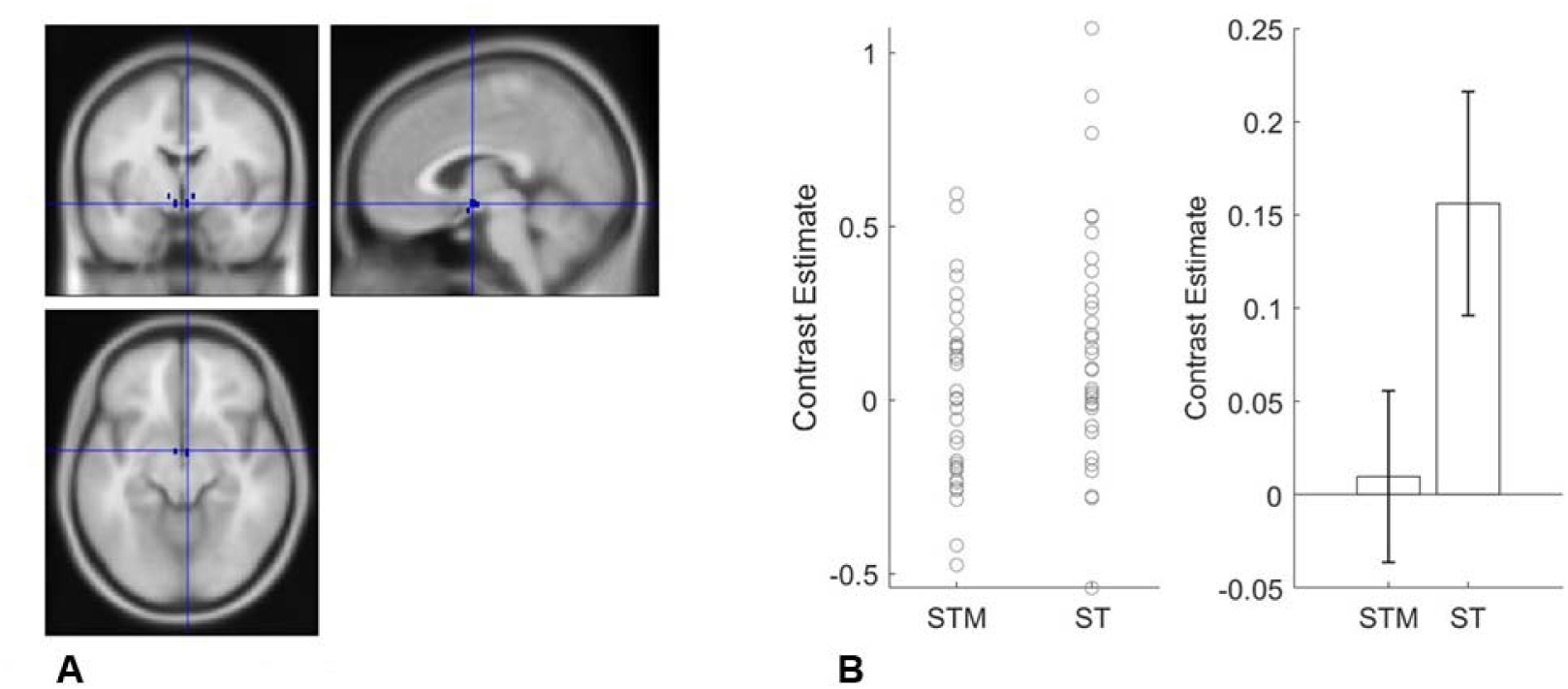
**A**. Hypothalamus ROI. **B.** Contrast estimates extracted from ROI using marsbar for stevia plus modifier (STM) and stevia (ST), (error bars, SEM).

**Figure 4.**
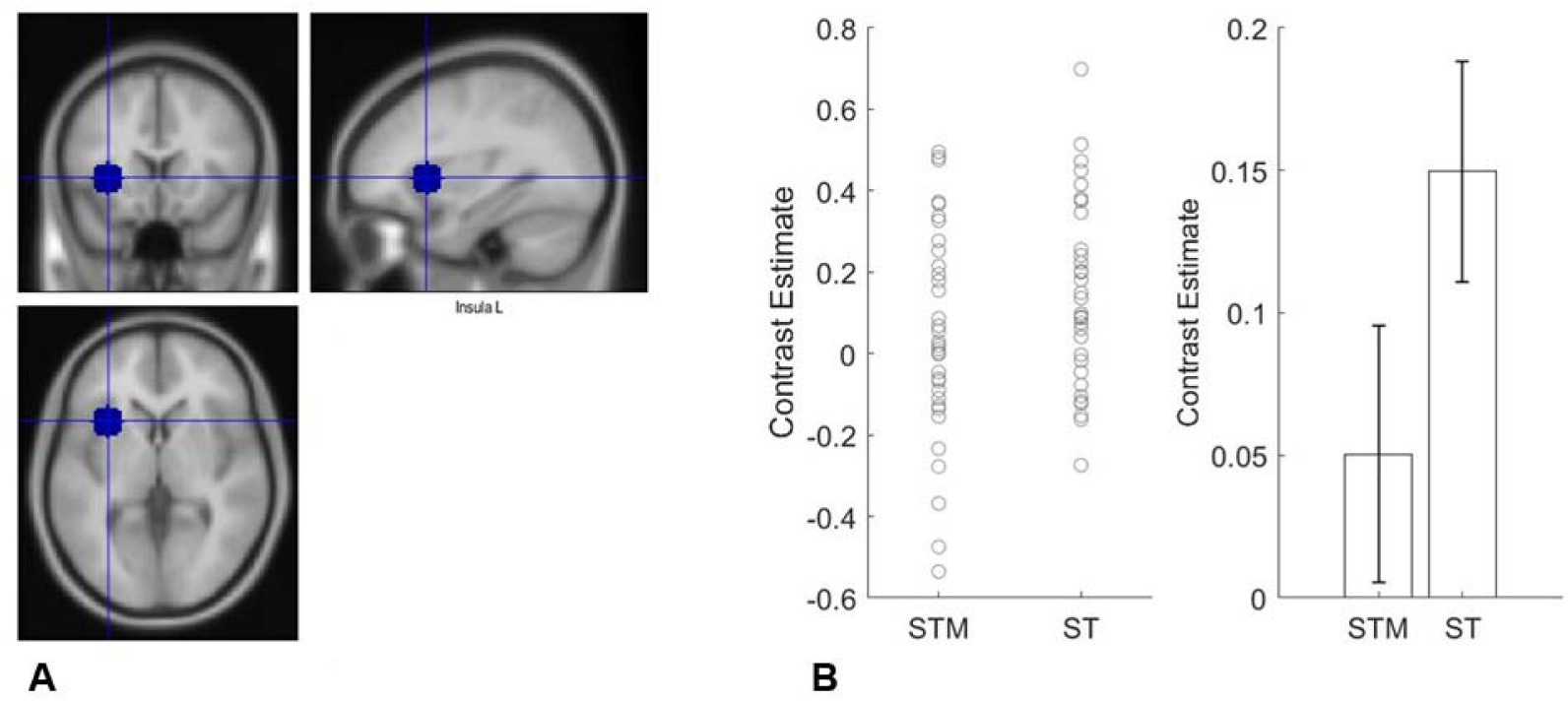
**A**. Left anterior insula ROI. **B.** Contrast estimates extracted from ROI using marsbar for stevia plus modifier (STM) and stevia (ST), (error bars, SEM).

#### S vs STM

We found increased neural activity in the S vs STM in the left anterior insula ROIs (t=2.69, p=0.005, Cohens d =0.46) and right anterior insula ROI (t=3.03, p=0.002, Cohens d =0.52) (Table S5).

#### S vs ST

We found increased neural activity in the S vs ST in the right postcentral gyrus ROI (t=2.26, p= 0.01, Cohens d =0.39) and right anterior insula ROI (t=1.99, p= 0.03, Cohens d =0.34) (Table S6). When correcting for multiple comparisons the postcentral gyrus activity remained significant (Figure 5).

**Figure 5.**
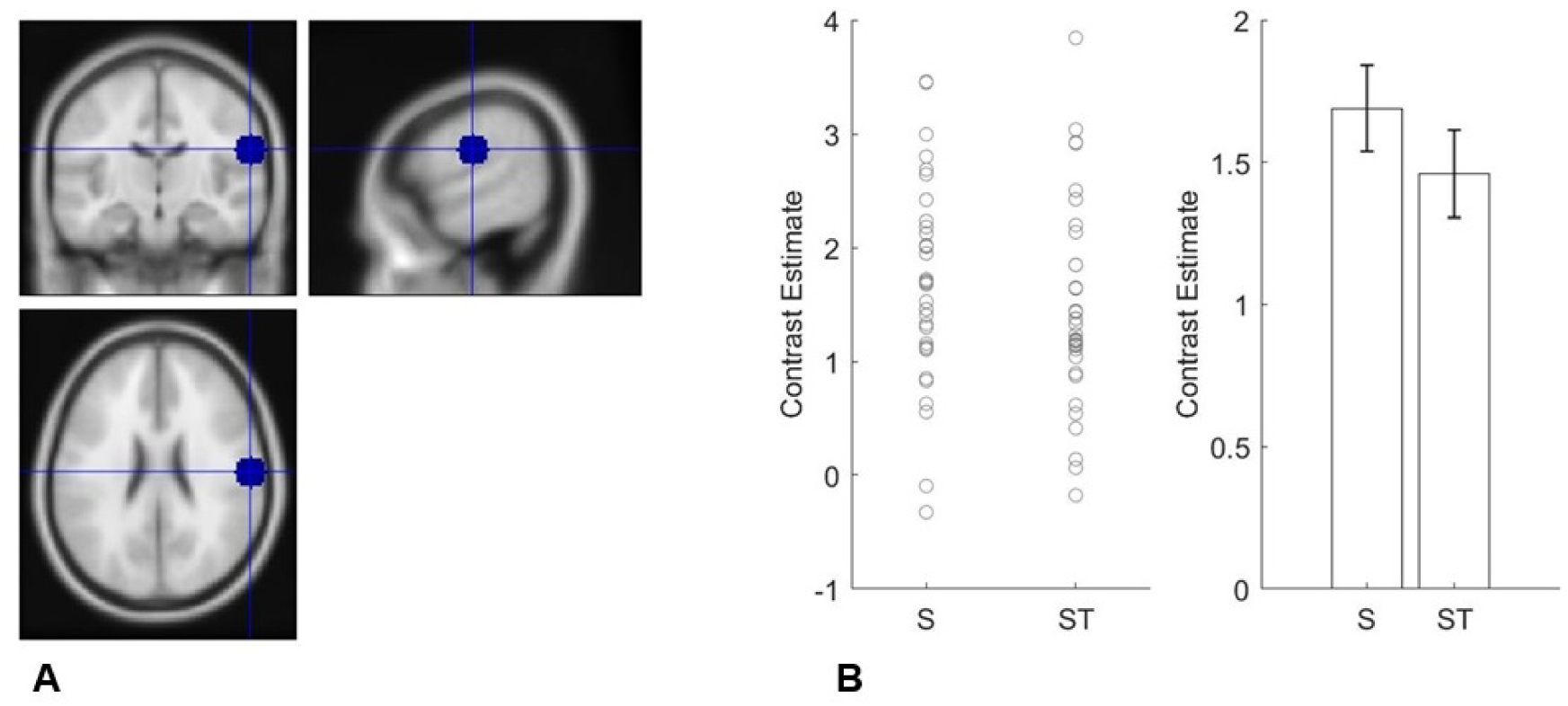
**A**. Right postcentral gyrus ROI. **B.** Contrast estimates extracted from ROI using marsbar for sucrose (S) and stevia (ST) (error bars, SEM).

### Temporal effects

#### STM vs ST

We also examined the time to peak activity after the taste for stevia stimuli. Using repeated measures ANOVA with ROI (11 levels) as a within subjects factor and condition as a second within subjects factor (ST, STM). We found a main effect of ROI (F=9.54 (5.94,196.15) p<0.001), no main effect of condition (F=1.12 (1,33) p=0.299), and no ROI * condition interaction (F=1.29 (6.94,229.0) p=0.258).

#### S vs ST

We also examined the time to peak activity after the taste for sucrose and stevia stimuli. Using repeated measures ANOVA with condition as a within subjects factor (sucrose, stevia) and ROI (postcentral, insula, NAcc, amygdala, hypothalamus) as a within subjects factor we found no main effect of conditon (F=0.001 (1,33) p=0.97), a main effect of ROI (F=6.81 (6.5,217.6) p<0.001), but no ROI * condition interaction (F=1.26 (6.3,210.6) p=0.27).

### Parametric modulation Pleasantness

We found as STM pleasantness increased activity in the left anterior insula (rho = −0.40, p = 0.009) and posterior insula (rho = −0.40, p = 0.01) decreased, while activity in the hypothalamus increased (rho = 0.34, p = 0.03).

We found as ST pleasantness increased activity in the left (rho = −0.41, p = 0.008) right anterior insula (rho = −0.42, p = 0.006) (Figure 6) and left posterior insula (rho = −0.56, p = 0.0006) decreased.

**Figure 6.**
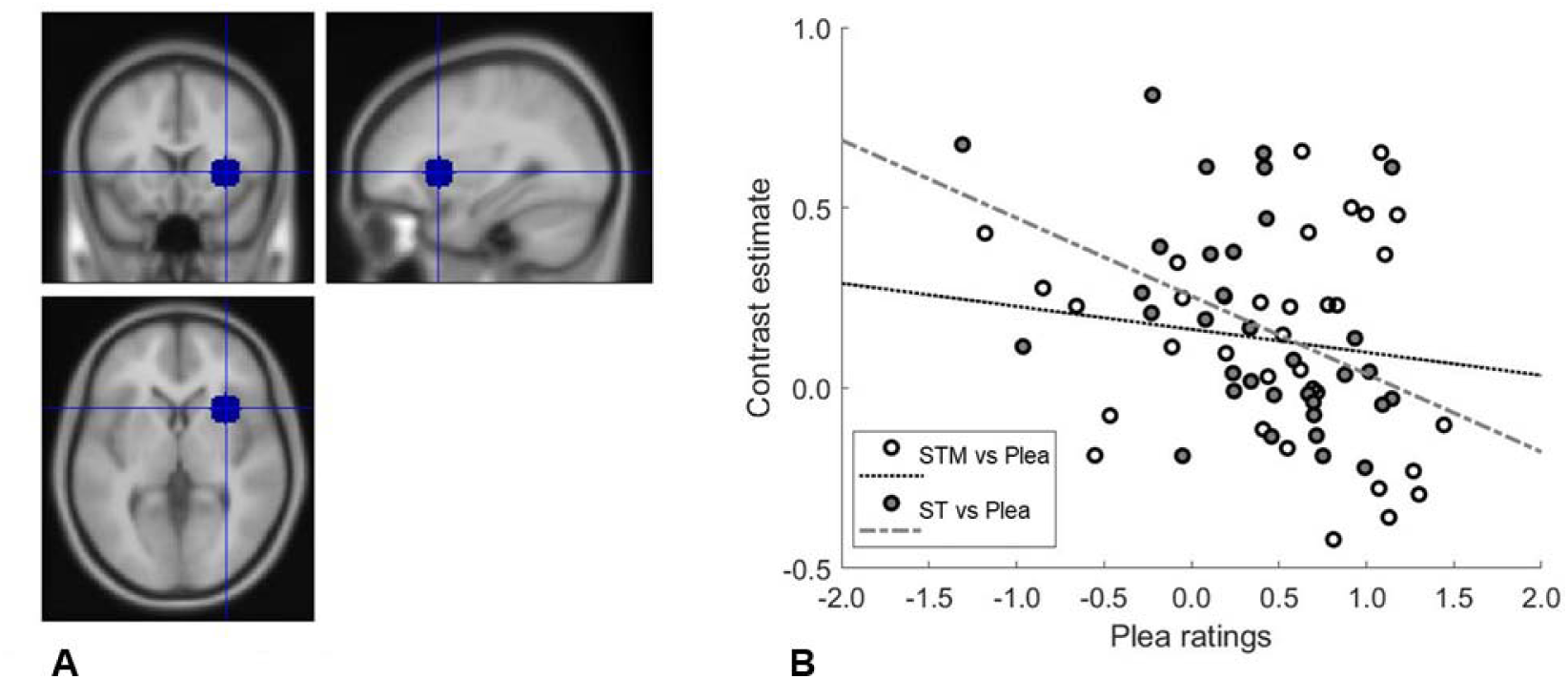
**A**. Right anterior insula ROI. **B**. Stevia (ST) pleasantness correlation with right anterior insula (rho=-0.422, p=0.006) and stevia plus modifier (STM) pleasantness correlation with right anterior insula (rho=-0.145, p=0.207) contrast estimates extracted from ROI using marsbar.

As S pleasantness increased activity in the posterior insula decreased (*rho* = −0.32, p = 0.03), however this did not survive multiple comparisons correction.

### Bitterness

As ST bitterness increased activity in the right amygdala increased (rho = 0.29, p = 0.05), but this did not survive corrections for multiple comparisons (Figure 7).

**Figure 7.**
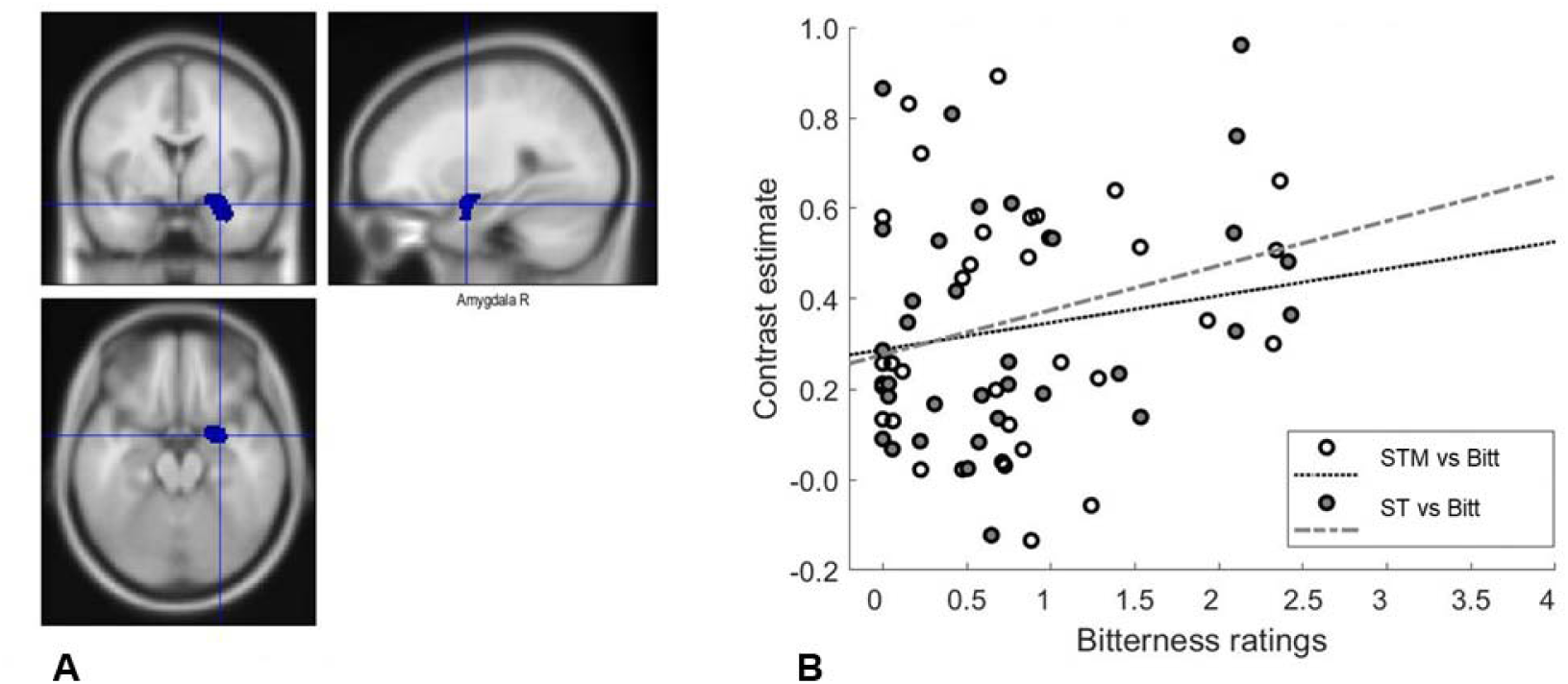
**A**. Right amygdala ROI. **B**. **B**. Stevia (ST) bitterness correlation (rho=0.291, p=0.05) and stevia plus modifier (STM) bitterness correlation (rho=0.157, p=0.187) contrast estimates extracted from ROI using marsbar.

### Mouth Fullness

As STM mouth fullness increased activity in the right anterior insula (rho =0.28, p=0.05) left NAcc (rho = 0.43, p = 0.01) and hypothalamus (rho = 0.36, p = 0.02) increased. As ST mouth fullness increased activity in the left NAcc (rho = 0.35, p = 0.03) increased and as S mouth fullness increased activity in the left NAcc (rho = 0.30, p = 0.05) increased. Only the correlation between the left NAcc and STM mouth fullness survived correction for multiple comparisons.

## Discussion

This is the first study to examine the effects of adding a FMP to the natural sweetener stevia in the human brain. We found synergistic neural activity in the postcentral gyri, parietal and occipital regions for stevia combined with the modifier compared to the sum of the activation to stevia alone and the modifier alone.

Reported in meta-analyses the postcentral gyri and parietal regions are responsive to sucrose (Roberts et al., 2020). The postcentral gyrus, part of the somatosensory cortex, is also influenced by sweet taste intensity (Small, 2012) (van Meer et al., 2023) while the parietal lobe supports sensory perception and integration, including taste and smell (Johns, 2014). The superior parietal lobe, closely connects to the occipital lobe, and thus contributes to attention and visuospatial processing (Johns, 2014). Therefore neural super-additivity in these regions could underpin the potential mechanisms by which non-nutrient sweeteners added to modifiers could be made more acceptable in diets, thus aiding sugar reduction in foods (Ko et al., 2025).

In the ROI analysis we found reduced activation of the hypothalamus to the stevia plus modifier vs stevia alone. The hypothalamus is regarded as one of the core processors in the control of appetite (Berthoud et al., 2017). Some studies report that non-nutrient sweeteners either have no effect on the hypothalamus (Smeets et al., 2005a) or a transient deactivation of the hypothalamus (Van Opstal, Kaal, et al., 2019). While others show that glucose ingestion results in decreased activity in the hypothalamus (Little et al., 2014; Simon et al., 2020; Smeets et al., 2005a, 2005b). Therefore, our finding could suggest that the FMP is producing neural effects like sucrose. Consistent with this, we did not find a significant neural difference when directly comparing hypothalamus activity between sucrose and the stevia plus modifier.

Next, we examined the neural differences in sucrose vs stevia given the paucity of data on this direct comparison. At the whole brain level, we found increased activity to sucrose in the insula and rolandic operculum, but only at an uncorrected threshold p<0.001. In the ROI analysis we found increased post central gyrus to sucrose vs stevia. The insula is central to taste processing and the postcentral gyrus to somatosensory processing (Yeung & Wong, 2020) (Small, 2012) while the rolandic operculum is involved in flavour perception and olfactory–gustatory integration (Seubert et al., 2015; Suen et al., 2021). Therefore, our results could reflect a more “objective” sensing of sweetness for sucrose vs stevia (van Meer et al., 2023) in line with studies reporting a physiological impact of sucrose vs stevia on EEG activity (Mouillot et al., 2020).

Finally, when examining the relationships between brain activity and the subjective experiences we found that the insula tracked the pleasantness of the stevia conditions more than the sucrose. As the stevia conditions were rated less pleasant than the sucrose this finding could reflect the insulas role in tracking unpleasant/other salient effects of stevia (Rolls, 2016). This is also supported by our finding that the amygdala, known for its role in both food aversion, salience processing (Izadi & Radahmadi, 2022) and the detection of bitterness (Hoogeveen et al., 2015) tracked the bitterness of the stevia but not the stevia plus modifier, despite no significant ratings of bitterness for stevia vs the stevia plus modifier.

We also found that the hypothalamus, a region that controls feeding behaviour (Van Opstal, van den Berg-Huysmans, et al., 2019) tracked the pleasantness of the stevia plus modifier condition but not the stevia. The hypothalamus, along with the NAcc (reward region) also tracked the mouth fullness of the stevia plus modifier condition, more so than the stevia.

Taken together, these findings suggest that adding a modifier to stevia could increase unconscious desirability for stevia outside of subjective awareness by masking its bitterness and increasing its mouth fullness. Future studies could examine modulation of neural regions such as the amygdala by FMPs can predict subsequent food consummation (Tiedemann et al., 2020).

## Acknowledgements

We would like to thank Dr Shan Shen and the staff at the Centre for Neuroscience and Neurodynamics (CINN) at the University of Reading for their help with the scanning.

## Author Contributions

CMcC, TE, JS, WS and HK conceived and designed research. JS, WS and TE prepared and supplied the study samples. HK and TE conducted research and analysed the data supported by CMcC. CMcC and HK drafted the article. All authors actively participated in editing and reviewing the manuscript.

## Data Accessibility Statement

The data that support the findings of this study are available from the corresponding author, [CMC], upon reasonable request.

