## Supplement for "Human Neural Synergy when combining Stevia with a Flavor Modifer and the Neural effects of Sucrose vs Stevia"

**Supplement doc**

**Whole brain analyses**

*Main effects of taste stimuli*

**Table** **S1:**

| **Sucrose - control**  Threshold: p=0.05 FWE corrected | | | | | | | |
| --- | --- | --- | --- | --- | --- | --- | --- |
| **Region** | **x** | **y** | **z** | **Z-score** | **voxels** | **p(FWE-corr)** | **p(FDR-corr)** |
| Postcentral gyrus | -42 | -18 | 52 | 6.31 | 152 | < 0.0001 | < 0.0001 |
| Precentral gyrus | -35 | -23 | 55 | 6.02 |  |  |  |
| Supplementary Motor Area | 4 | 10 | 55 | 6.05 | 252 | <0.0001 | < 0.0001 |
| Insula | 37 | 20 | 7 | 5.70 | 106 | < 0.0001 | < 0.0001 |
| Mid Cingulate gyrus | -8 | 25 | 31 | 5.46 | 10 | <0.001 | =0.05 |
| Hippocampus | 42 | -30 | -8 | 5.40 | 22 | < 0.0001 | =0.004 |
| Insula | -35 | 15 | 7 | 5.26 | 29 | < 0.0001 | =0.001 |
| Caudate | 18 | 15 | 7 | 5.13 | 23 | < 0.0001 | =0.004 |
| Insula/FO | -42 | 18 | 0 | 5.13 | 10 | 0.0008 | 0.05 |
| FO, Frontal operculum, Results remain the same with gender, hunger level and scan time added as covariates | | | | | | | |

**Table S2:**

| **Stevia - control**  Threshold: p=0.05 FWE corrected | | | | | | | |
| --- | --- | --- | --- | --- | --- | --- | --- |
| **Region** | **x** | **y** | **z** | **Z-score** | **voxels** | **p(FWE-corr)** | **p(FDR-corr)** |
| Precentral gyrus | -35 | -28 | 64 | 6.48 | 196 | < 0.0001 | < 0.0001 |
| Postcentral gyrus | -40 | -21 | 52 | 6.17 |  |  |  |
| Caudate | 13 | 15 | 4 | 6.14 | 152 | < 0.0001 | < 0.0001 |
| Caudate | -8 | 10 | 2 | 5.38 |  |  |  |
| Putamen | -18 | 15 | 0 | 5.07 |  |  |  |
| Insula | -30 | 25 | 7 | 5.99 | 78 | < 0.0001 | < 0.0001 |
| Insula | 30 | 27 | 2 | 5.89 | 72 | < 0.0001 | < 0.0001 |
| Supplementary Motor Area | -4 | 15 | 45 | 5.63 | 26 | < 0.0001 | =0.0006 |
| Supplementary Motor Area | 4 | 3 | 60 | 5.37 | 46 | < 0.0001 | < 0.0001 |
| Results remain the same with gender, hunger level and scan time added as covariates | | | | | | | |

**Table S3:**

| **Modifier - control**    Threshold: p=0.05 FWE corrected | | | | | | | |
| --- | --- | --- | --- | --- | --- | --- | --- |
| **Region** | **x** | **y** | **z** | **Z-score** | **voxels** | **p(FWE-corr)** | **p(FDR-corr)** |
| Insula | -30 | 25 | 2 | 6.34 | 86 | < 0.0001 | < 0.0001 |
| Insula/FO | -40 | 13 | 2 | 5.26 |  |  |  |
| Caudate | -11 | 10 | 7 | 6.25 | 170 | < 0.0001 | < 0.0001 |
| Insula | 30 | 27 | 0 | 6.17 | 44 | < 0.0001 | < 0.0001 |
| Supplementary Motor Area | 4 | 8 | 55 | 5.98 | 269 | <0.0001 | < 0.0001 |
| Precentral gyrus | -42 | -18 | 55 | 5.49 | 50 | < 0.0001 | < 0.0001 |
| Insula/FO | 35 | 15 | 9 | 5.46 | 23 | < 0.0001 | =0.002 |
| Note: FO= Frontal operculum. Results remain the same with gender, hunger level and scan time added as covariates | | | | | | | |

| **Table S4: ROI Analysis ST vs STM** | | | | | | |
| --- | --- | --- | --- | --- | --- | --- |
| **Region** | **ST**  **(Mean**  ± **SD)** | **STM**  **(Mean** ± **SD)** | **P value *(cohens d)*** | **ST**  **(Mean** ± **SD)** | **STM**  **(Mean** ± **SD)** | **P value**  ***(cohens d)*** |
|  | **Left** | | | **Right** | | |
| Postcentral gyrus | 1.31  ± 0.67 | 1.24  ± 0.70 | 0.25  (0.11) | 1.46  ± 0.89 | 1.39  ± 0.82 | 0.31  (0.08) |
| Insula (anterior) | 0.15  ± 0.23 | 0.05  ± 0.26 | 0.02  (0.36) | 0.18  ± 0.29 | 0.13  ± 0.29 | 0.15  (0.18) |
| Insula (posterior) | 0.19  ± 0.21 | 0.17  ± 0.19 | 0.36  (0.06) | 0.21  ± 0.32 | 0.22  ± 0.23 | 0.42  (-0.03) |
| NAcc | 0.20  ± 0.35 | 0.10  ± 0.29 | 0.06  (0.29) | 0.14  ± 0.33 | 0.16  ± 0.29 | 0.34  (-0.07) |
| Amygdala | 0.32  ± 0.32 | 0.26  ± 0.29 | 0.11  (0.21) | 0.36  ± 0.26 | 0.33  ± 0.26 | 0.35  (0.06) |
| Hypothalamus | 0.16  ± 0.34 | 0.01  ± 0.26 | 0.008* (0.43) |  |  |  |
| Mean, contrast estimates extracted in each ROI. * Survives multiple comparison correction (0.05/5 ROIs,p=0.01) | | | |  |  |  |

| **Table S5: ROI Analysis S vs STM** | | | | | | |
| --- | --- | --- | --- | --- | --- | --- |
| **Region** | **S**  **(Mean ± SD)** | **STM**  **(Mean ± SD)** | **P value**  ***(cohens d)*** | **S**  **(Mean ± SD)** | **STM**  **(Mean ± SD)** | **P value**  ***(cohens d)*** |
|  | **Left** | | | **Right** | | |
| Postcentral gyrus | 1.44  ± 0.80 | 1.24  ± 0.70 | 0.02  (0.36) | 1.69  ± 0.88 | 1.39  ± 0.82 | 0.02  (0.37) |
| Insula (anterior) | 0.21  ± 0.32 | 0.05  ± 0.26 | 0.005*  (0.46) | 0.26  ± 0.32 | 0.13  ± 0.29 | 0.002*  (0.52) |
| Insula (posterior) | 0.17  ± 0.31 | 0.17  ± 0.19 | 0.48  (-0.01) | 0.30  ± 0.28 | 0.22  ± 0.23 | 0.06  (0.28) |
| NAcc | 0.24  ± 0.32 | 0.10  ± 0.29 | 0.02  (0.40) | 0.22  ± 0.34 | 0.16  ± 0.29 | 0.20  (0.15) |
| Amygdala | 0.33  ± 0.26 | 0.26  ± 0.29 | 0.07  (0.26) | 0.38  ± 0.25 | 0.33  ± 0.26 | 0.20  (0.15) |
| Hypothalamus | 0.12  ± 0.24 | 0.01  ± 0.26 | 0.02  (0.38) |  | | |
| Mean, contrast estimates extracted in each ROI. * Survives multiple comparison correction (0.05/5 ROIs, p=0.01) | | | |  | | |

| **Table S6: ROI Analysis S vs ST** | | | | | | |
| --- | --- | --- | --- | --- | --- | --- |
| **Region** | **S**  **(Mean** ± **SD)** | **ST**  **(Mean**  ± **SD)** | **P value *(cohens d)*** | **S**  **(Mean** ± **SD)** | **ST**  **(Mean**  ± **SD)** | **P value *(cohens d)*** |
|  | **Left** | | | **Right** | | |
| Postcentral gyrus | 1.44  ±0.80 | 1.31  ± 0.67 | 0.07  (0.25) | 1.69  ±0.88 | 1.46  ± 0.89 | 0.01*  (0.39) |
| Insula (anterior) | 0.21  ±0.32 | 0.15  ± 0.23 | 0.09  (0.24) | 0.26  ±0.32 | 0.18  ± 0.29 | 0.03  (0.34) |
| Insula (posterior) | 0.17  ±0.31 | 0.19  ± 0.21 | 0.37  (-0.06) | 0.30  ±0.28 | 0.21  ± 0.32 | 0.05  (0.28) |
| NAcc | 0.24  ±0.32 | 0.20  ± 0.35 | 0.30  (0.10) | 0.22  ±0.34 | 0.14  ± 0.33 | 0.09  (0.24) |
| Amygdala | 0.33  ±0.26 | 0.32  ± 0.32 | 0.41  (0.04) | 0.38  ±0.25 | 0.36  ± 0.26 | 0.31  (0.09) |
| Hypothalamus | 0.12  ±0.24 | 0.16  ± 0.34 | 0.27  (-0.11) |  |  |  |
| Mean, contrast estimates extracted in each ROI. * Survives multiple comparison correction (0.05/5 ROIs,p=0.01) | | | |  |  |  |
